# Development of Inflammatory Bowel Disease is Linked to a Longitudinal Restructuring of the Gut Metagenome in Mice

**DOI:** 10.1101/173823

**Authors:** Thomas Sharpton, Svetlana Lyalina, Julie Luong, Joey Pham, Emily M. Deal, Courtney Armour, Christopher Gaulke, Shomyseh Sanjabi, Katherine S. Pollard

## Abstract

The gut microbiome is linked to inflammatory bowel disease (IBD) severity and altered in late stage disease. However, it is unclear how gut microbial communities change over the course of IBD development, especially in regards to function. To investigate microbiome mediated disease mechanisms and discover early biomarkers of IBD, we conducted a longitudinal metagenomic investigation in an established mouse model of IBD, where dampened TGF-β signaling in T cells leads to peripheral immune activation, weight loss, and severe colitis. IBD development is associated with abnormal gut microbiome temporal dynamics, including dampened acquisition of functional diversity and significant differences in abundance trajectories for KEGG modules such as glycosaminoglycan degradation, cellular chemotaxis, and type III and IV secretion systems. Most differences between sick and control mice emerge when mice begin to lose weight and heightened T cell activation is detected in peripheral blood. However, lipooligosaccharide transporter abundance diverges prior to immune activation, indicating that it could be a pre-disease indicator or microbiome-mediated disease mechanism. Taxonomic structure of the gut microbiome also significantly changes in association with IBD development, and the abundance of particular taxa, including several species of *Bacteroides*, correlate with immune activation. These discoveries were enabled by our use of generalized linear mixed effects models to test for differences in longitudinal profiles between healthy and diseased mice while accounting for the distributions of taxon and gene counts in metagenomic data. These findings demonstrate that longitudinal metagenomics is useful for discovering potential mechanisms through which the gut microbiome becomes altered in IBD.

**Importance:** IBD patients harbor distinct microbial communities with different functional capabilities compared to healthy people. But is this cause or effect? Answering this question requires data on changes in gut microbial communities leading up to disease onset. By performing weekly metagenomic sequencing and mixed effects modeling on an established mouse model of IBD, we identified several functional pathways encoded by the gut microbiome that covary with host immune status. These pathways are novel early biomarkers that may either enable microbes to live inside an inflamed gut or contribute to immune activation in IBD mice. Future work will validate the potential roles of these microbial pathways in host-microbe interactions and human disease. This study is novel in its longitudinal design and focus on microbial pathways, which provided new mechanistic insights into the role of gut microbes in IBD development.

## Background

Inflammatory bowel disease (IBD) is an increasingly prevalent chronic autoimmune disease wherein the cells of the immune system attack intestinal tissue (1-3). Quality of life deteriorates, and patients die in severe cases. Unfortunately, the etiology of disease remains unclear and is likely complex (4). Discovery of the factors that contribute to IBD onset, development, and severity is needed to ensure accurate and effective health care. Epidemiological studies and animal model experiments have identified genetic (5-7) and lifestyle factors that associate with IBD, including diet (8) and exercise (9). But these factors are not precise predictors of disease risk, severity, or response to treatment, and many questions remain regarding disease mechanisms. Elucidating the cryptic etiology of IBD would enable new preventative measures, diagnostics, and therapies.

Recent work has implicated the gut microbiome in the development and severity of IBD (10). Individuals afflicted with Crohn’s disease or ulcerative colitis, the two principal clinical forms of IBD, harbor distinct taxa relative to healthy controls (11-14). Shotgun metagenomics further revealed that the abundance of several microbial metabolic pathways are significantly altered in IBD guts (13, 15, 16). These associations may be causal, because gut microbes can influence the immune system and intestinal homeostasis. For example, immunosuppressive regulatory T cells (Tregs) are prevalent in the colonic lamina propria (LP) compared to other organs. But, their numbers are reduced in germ-free or antibiotic-treated mice, suggesting that microbiota affect colonic differentiation of peripheral Tregs (pTregs) (17, 18). A similar loss of Tregs occurs in people with polymorphisms in IBD-susceptibility genes that promote defects in Treg responses (19). Thus, gut microbes have the potential to interact with immune cells and this interaction can be altered due to host genetics and other risk factors in the development of IBD.

We hypothesized that the changes in immune status of individuals with IBD are associated with temporal alterations in the functional capabilities of their gut microbiota. Understanding how the gut microbiome dynamically changes during IBD and how these changes relate to host symptoms and immune activation could clarify which microbiomic alterations contribute to disease onset and progression and which alterations respond to disease. We are particularly interested in elucidating specific microbial pathways that may induce or exacerbate immune activation and distinguishing these from pathways required for survival in an inflamed intestinal environment. Addressing these questions requires a prospective, longitudinal study of the microbiome in IBD.

Longitudinal investigations of the microbiome have tended to focus on taxonomic rather than functional changes (20, 21). One study used 16S sequencing in the T-bet-/-RAG2-/-Ulcerative Colitis (TRUC) mouse model of inflammatory disease to identify how gut microbiome taxonomic composition changes over the course of treatment-induced remission and then imputed how microbial pathway abundances might change over time with ancestral state reconstruction techniques (22). Shotgun metagenomic sequencing provides direct insight into the functions encoded in the microbiome, but it has not been applied to a longitudinal investigation of IBD. As a result, our insight into how the gut microbiome operates dynamically in association with disease development is limited.

Mouse models of disease present an opportunity to quantify the longitudinal covariation between gut microbiome functions and IBD development while overcoming the challenges associated with a prospective human study and reducing the extensive genetic, lifestyle, and microbiome variation among humans. We implemented this approach using a well-documented IBD model (23-29), where TGF-β dominant negative receptor II is driven by the CD4 promoter (CD4-dnTβRII) (30), called DNR hereafter. TGF-β is important for inducing pTreg differentiation (31), and its signaling in naive T cells results in activation and nuclear translocation of Smad2/3 molecules and regulation of target genes, including Foxp3 (32-34). Foxp3 then provides a positive feedback loop by downregulating Smad7, thereby reducing its inhibition of TGF-β signaling (35). Absence of TGF-β signaling in T cells results in loss of Foxp3 expression and defective *in vivo* expansion and immunosuppressive capacity of pTregs (36, 37). However, excess inflammation can also potently inhibit Foxp3 induction by TGF-β (38, 39), and the presence of certain inflammatory cytokines can instead divert differentiation of Tregs into pathogenic Th17 cells (40-45). Thus, due to TGF-β’s involvement in Treg cell differentiation, and the requirement for Treg produced IL-10 to maintain intestinal homeostasis, TGF-β signaling in T cells is an important component of intestinal immunity (46-54). Furthermore, mutations in both TGF-β and IL-10 signaling pathways have been implicated in human IBD (55-58). As a result of the blockage of TFGβ signaling on their T cells, and reduced number of pTregs, DNR animals develop spontaneous colonic inflammation and IBD that is akin to Crohn’s disease (30, 59). In addition to these physiological similarities, the DNR line serves as an effective model of human IBD because (i) human IBD is associated with mutations in SMAD3 (5, 60-62), a direct downstream target of TGFβ RII required for Foxp3 induction in the gut (33), and (ii) DNR mice model the documented effect of Smad7 overexpression in human IBD (63-65).

To obtain insight into how the longitudinal dynamics of the microbiome associate with IBD onset and progression, we followed DNR and littermate wild-type (hereafter, WT) controls from weaning through severe disease. We used shotgun metagenomics to quantify how fecal microbiome structure and function change over the course of disease development in DNR mice and identified components of the microbiome that both associate with and predict immune status. We focus on longitudinal changes in biological pathways (i.e., groups of genes performing a coherent function), using estimated abundances of KEGG modules from DNR and WT metagenomes. Our work indicates that the microbiome may contain biomarkers of IBD development, clarifies mechanisms through which the microbiome may contribute to disease development, and reveals how gut microbes operate to succeed in an inflamed intestinal environment.

## Results

Age-matched female WT and DNR littermates were monitored longitudinally for IBD development over a period of 9 weeks, starting at 4 weeks of age upon being weaned from their mother. As this is a T cell-mediated IBD model, we quantified peripheral CD4 and CD8 T cell activation by flow cytometry and measured the longitudinal change in the CD44^hi^ activated fraction, which includes both effector and memory T cells (Additional File 1: Figure S1). We also measured the weight of the animals over time (Fig. 1A). As expected, WT mice gained weight and maintained a constant fraction of activated T cells. DNR mice, conversely, stopped gaining weight and experienced a sharp increase in CD4 T cell activation followed by gradual increase in CD8 T cell activation starting at 7 weeks of age (Fig. 1). These results indicate that in our facility, the DNR mice develop signs of IBD starting around week 7 and full disease by week 9. DNR mice had to be euthanized by week 15, as they had lost more than 15% of their maximum body weight. Similar to the T cell activation phenotype observed in the blood after week 7 (Fig. 1B-C), the DNR animals had a larger fraction of activated T cells in the spleen and the gut-draining mesenteric lymph node (MLN) at week 15 (Additional File 2: Figure S2).

**Figure 1.**
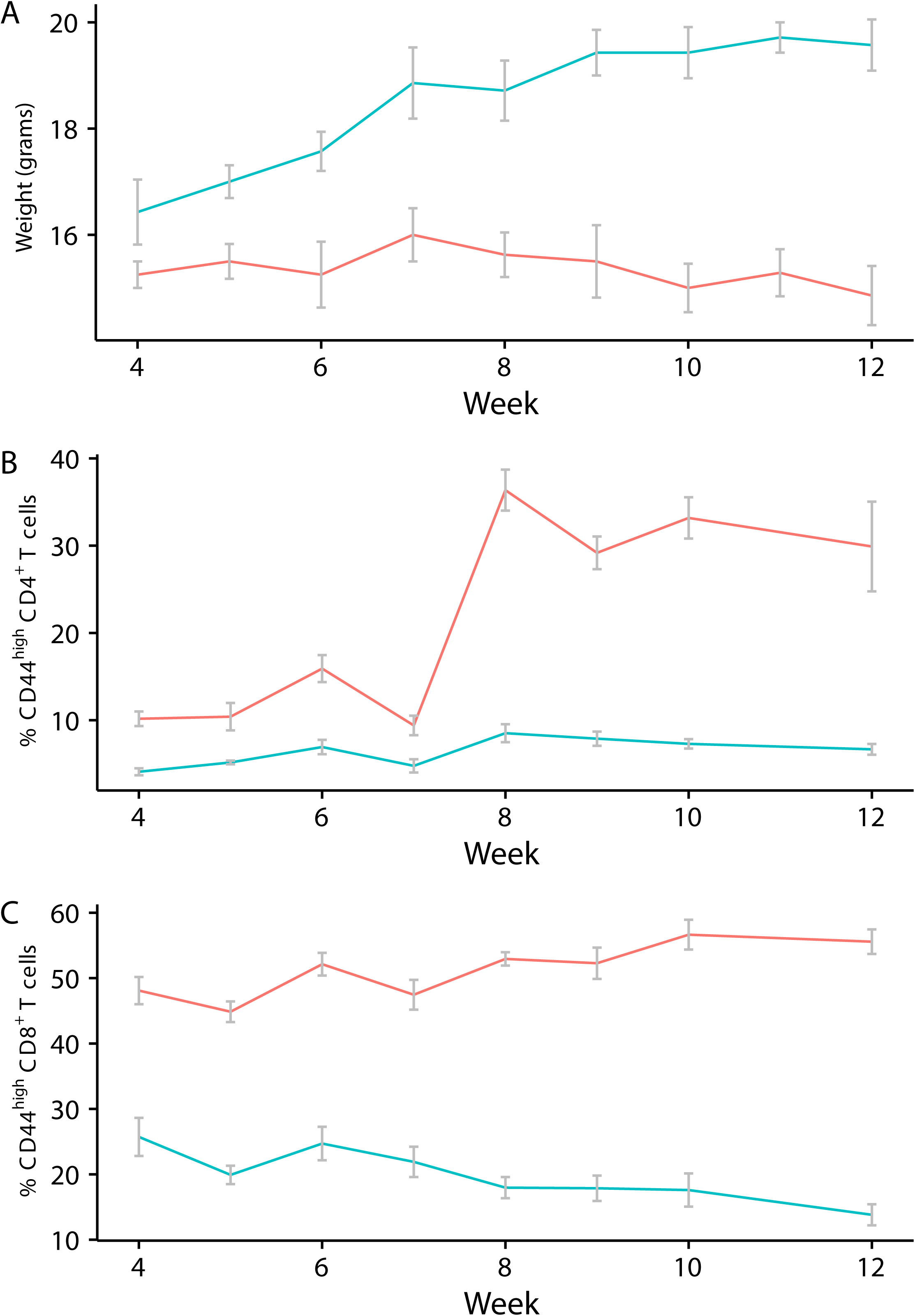
IBD development correlates with peripheral T cell activation in DNR mice. (**A**) Animal weight over time. N= 7 WT and 8 DNR mice. (**B**) Percent of activated CD4 T cells among peripheral blood mono-nuclear cells (PBMCs). (**C**) Percent of activated CD8 T cells among PBMCs. (**B-C**) N=6 WT and 6 DNR mice.

We used shotgun metagenomics to assess how the functional potential of the gut microbiome diversifies over the course of disease progression. Specifically, we collected stool samples from parallel cohorts of DNR and WT mice weekly and performed shotgun metagenomic sequencing from samples obtained at 4, 5, 6, 8, 10, 12, and 13 weeks of age (Additional File 3: Table S1). We then quantified the abundance of KEGG modules encoded in each metagenome with ShotMAP (16), which revealed 373 modules present in at least one sample. These module abundances were then used to quantify how the within-sample (alpha) diversity of microbiome functions varies over time in DNR and WT mice. A Kruskal-Wallis test of the change in KEGG module Shannon entropy over time (Additional File 4: Figure S3) found that the DNR mice are relatively stable in their functional alpha-diversity (p=0.47) as compared to WT mice (p=0.078). We also observed that functional alpha-diversity varies among individuals within a line over time, and that this variation differs between lines in association with disease activation (week 7). Specifically, the coefficient of variation of KEGG module Shannon entropy (CV) from a generalized linear model is higher among WT than DNR mice after disease activation (p = 0.0085). We also find that the CV is higher among DNR mice prior to activation, though this difference is reduced when the disproportionately variable week 5 samples are removed from the analysis (p=0.21). These results show that the functional diversity of the mouse gut microbiome is relatively constrained early in life but increases over the lifetimes of WT but not DNR individuals.

We then investigated how the composition of gut microbiome functions varies over time and between cohorts (DNR vs. WT) by using an abundance-weighted beta-diversity metric (Bray-Curtis dissimilarity). At a global level, KEGG module abundances were similar between DNR and WT mice prior to week 6, but then diverged over time as IBD developed in the DNR mice (Fig. 2). Furthermore, the diversity of KEGG modules found in a metagenome was significantly associated with the week that the sample was collected within the cohort (PERMANOVA p=0.01, R^2^=0.42), as well as cohort’s weekly mean activated T cell status (pcCD4tCD44hi, PERMANOVA p=0.001, R^2^=0.16). Thus, there exist microbiome-encoded functional modules that differ in abundance in association with IBD progression in DNR mice.

**Figure 2.**
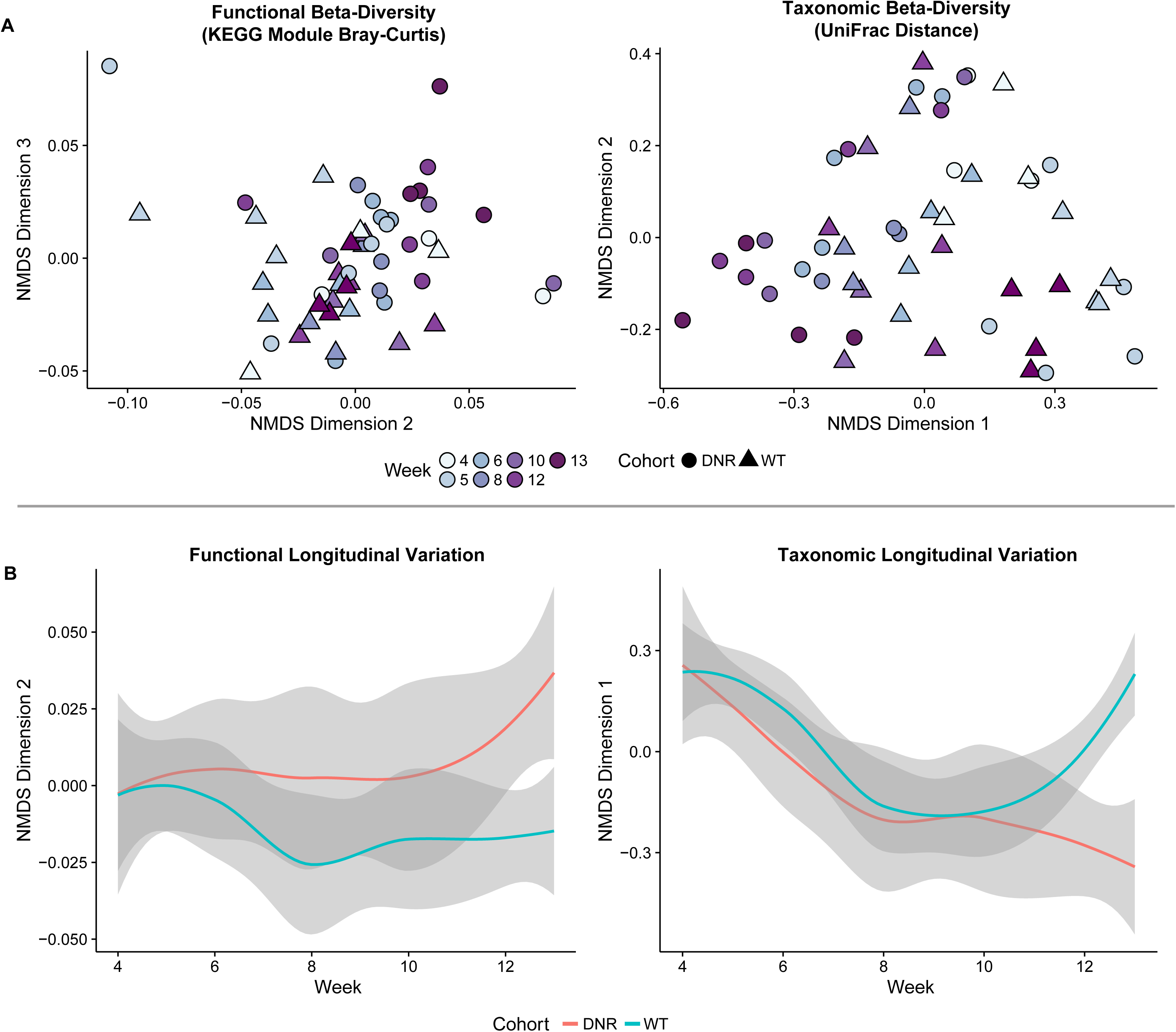
The taxonomic and functional diversity of the gut microbiome associates with IBD development. (A) NMDS ordination plots of the functional (left) and taxonomic (right) beta-diversity of samples from each line illustrate the significant divergence in beta-diversity between lines over time. Functional beta-diversity was measured as the Bray-Curtis dissimilarity based on KEGG module abundances, while taxonomic beta-diversity is the UniFrac distance of taxa detected in metagenomes. (B) The longitudinal variation of samples along selected NMDS dimensions similarly reveals how DNR and WT lines significantly diverge over time both in terms of their functional (left) and taxonomic (right) beta-diversity. Plotted is the smoothed LOESS trajectory of samples from each line over time, where grey areas represent 95% confidence intervals.

This temporal divergence in DNR versus WT microbiome functions is mirrored in the taxonomic structure of the microbiome (Fig. 2). The composition of the gut metagenomes is relatively similar between WT and DNR lines at early time points and begins to diverge at week 6. Additionally, the microbiomes of WT mice remain relatively consistent over time as compared to DNR mice, though they are not without temporal variation. Indeed, similar to the functional diversity analysis, the taxonomic beta-diversity of the microbiome significantly differs between the lines over time (PERMANOVA p=0.004, R^2^=0.46), though not with mean activated T cell status (PERMANOVA p=0.118, R^2^=0.046). Collectively, these analyses indicate that (1) the diversity and structure of the gut microbiome varies over time between 4 and 15 weeks of age in both WT and DNR mice, (2) WT and DNR microbiomes are generally consistent prior to immune activation in DNR mice, but diverge afterwards, and (3) immune activation is associated with changes in the subsequent succession of the gut microbiome.

Based on these observations, we assessed how specific components of the microbiome associate with disease development. A key novelty of our approach is the use of Tweedie compound Poisson generalized linear mixed effects models (GLMMs). These models allow us to test for differences in temporal trends in KEGG module abundance between DNR and WT mice while accounting for baseline differences between mice and genotypes, as well as DNA extraction kit effects. GLMMs enable accurate modeling of non-normally distributed abundance data and correctly account for multiple sources of variation (66), including the inter-subject variation that is present in repeated measures designs such as the longitudinal sampling of individual mice in our study. The Tweedie compound Poisson distribution, which is a weighted mixture between Poisson and Gamma distributions, has a number of other attractive features. Its exponential relationship between variance and mean captures the overdispersion that is frequently present in environmental DNA sequence data, and its point mass at zero allows for one-step fitting of zero-inflated data (versus fitting a model to determine feature presence/absence before modeling non-zero components, as in hurdle models). Additionally, the Tweedie compound Poisson is a continuous distribution, allowing us to use a normalized abundance measure as the dependent variable, instead of raw counts. We provide a more detailed description of the models used in our analysis in Additional Data File 5: Text S1.

We first looked at overall trends of abundance trajectories for DNR versus WT mice as quantified by the interaction between genotype and time in the GLMM. These analyses revealed 29 KEGG modules with significant differences in abundance trends between DNR and WT mice (FDR<0.05). The interaction coefficient was positive for 26 of the significant modules (Additional File 6: Table S2), which indicates that these modules became increasingly abundant in DNR versus WT mice over time. This set includes modules associated with uridine monophosphate biosynthesis (M00051), keratin sulfate degradation (M00079), and the type III secretion system (M00332). The three modules with negative interaction coefficients, indicating decreasing abundance in DNR versus WT mice over time (Fig. 3), are lysine biosynthesis (M00031), lipooligosaccharide transport (M00252), and melatonin biosynthesis (M00037).

**Figure 3.**
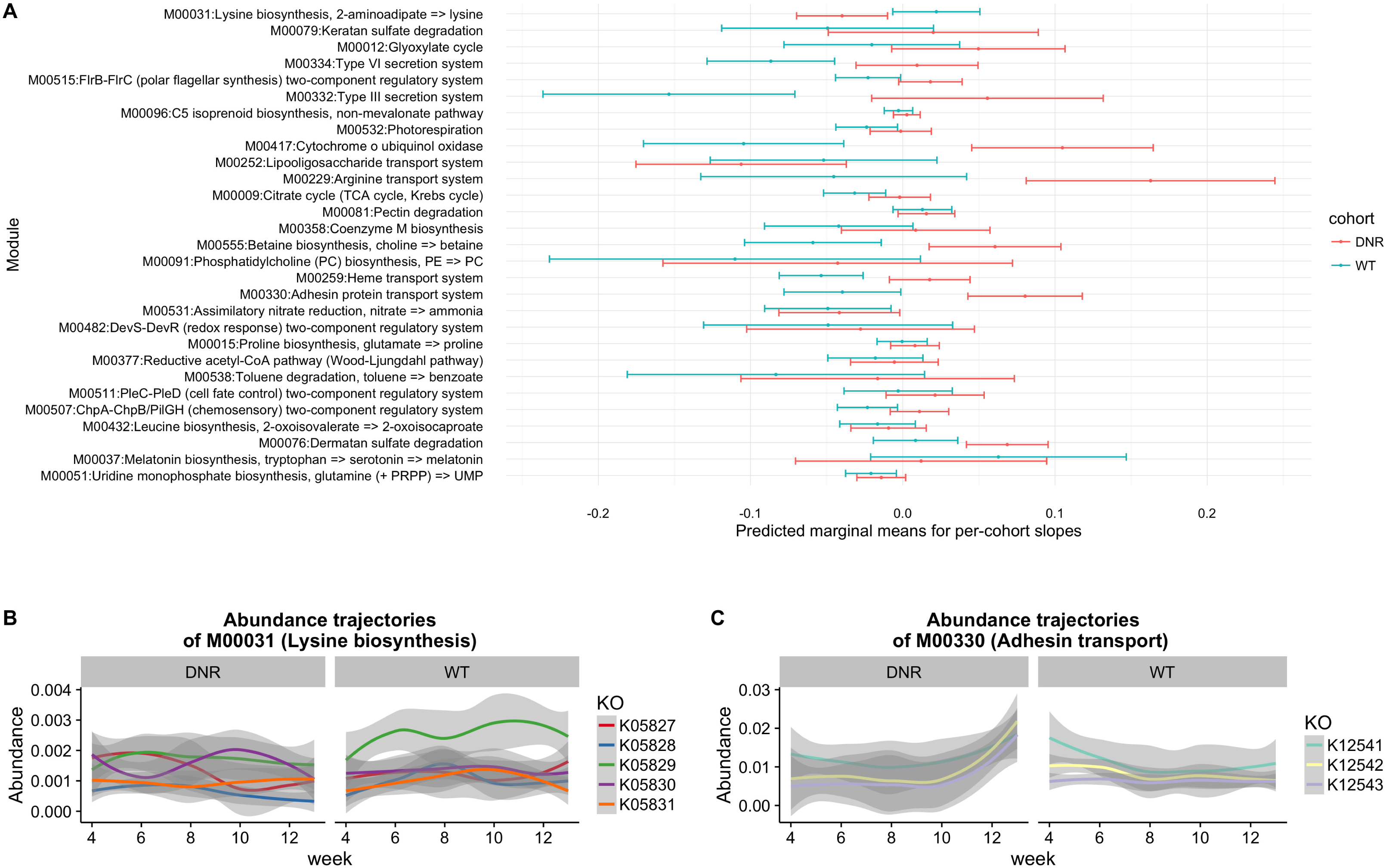
Summary of GLMM results of 29 modules with significant time by group interaction. (A) The quantity plotted is the predicted marginal mean (PMM) of the slope coefficients. Significance testing was done by comparing goodness of fit of full and reduced GLMM specifications, and the full model was used to produce the PMM estimates shown here. This quantity was primarily calculated to get a succinct summary of the direction of temporal change, and does not always coincide with the interaction coefficient that is the focus of the main analysis. The estimates were obtained by running the lstrends function from the lsmeans R package (136) (B) The underlying KO abundance trajectories of a significant module (M00031: Lysine biosynthesis) that decreases in DNR mice and increases in WT mice over time, as evidenced by a negative and positive model slope, respectively. (C) A similar plot as in (B), except that this significant module (M00330: Adhesin transport) significantly increases in abundance over time in DNR mice, while it does not change in abundance in WT mice. For both (B) and (C) the shaded ribbons represent LOESS confidence bounds.

To obtain improved temporal resolution regarding the divergence of module abundance in DNR mice, we extended our GLMMs to include a “hinge” at week 7, which is when immune activation initiates in DNR mice. This segmented regression approach has the potential to discover modules that diverge in abundance between DNR and WT mice either between weeks 4 and 7 or between weeks 7 and 13. Only 13 of the 29 previously identified modules exhibited a significant effect when using segmented regression (Fig. 4), likely due to a loss of power from partitioning the data into two smaller sets of samples. However for these 13 modules, our results clarify when DNR and WT abundances began to diverge (Additional File 7: Table S3). The predominant pattern was similar module abundance prior to week 7, followed by divergence after immune activation (11/13 modules). This pattern suggests that these modules respond to disease or play a role in disease progression.

**Figure 4.**
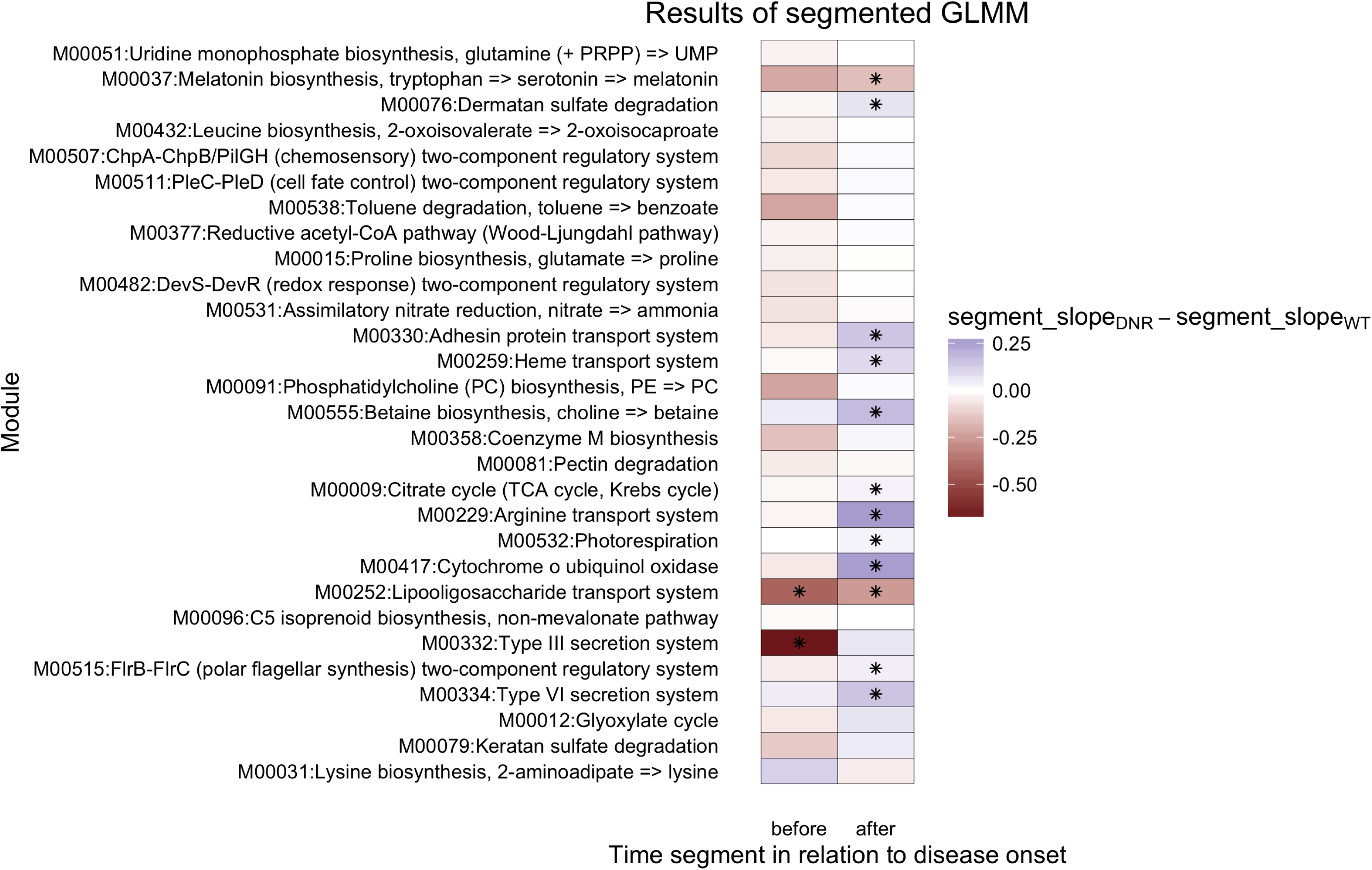
Modules with significantly differing slopes between groups show primarily post-disease-onset differences when analyzed with a segmented GLMM. For each cohort, the segmented GLMM estimates two separate WT slopes (pre-week7 and post-week7) and two deviations from those slopes, which represent the time by group interaction that measures how DNR slopes differ from WT slopes. Plotted here are the estimates of these deviations, with asterisks marking coefficients that were significantly non-zero with B-H corrected p-value of <0.2

Lipooligosaccharide transport (M00252), which is a two-component system with an unknown substrate in the mammalian gut, was the only module that stratified DNR and WT mice both before and after disease onset. To further investigate the potential taxa that may drive this particular signal, we assessed the taxonomic source of the KEGG sequences that recruited metagenomic reads into the module. We also quantified the distance covariance (67) between the longitudinal trajectories of the KEGG Orthology Groups (KOs) that comprise the module and each observed species’ trajectory. The result was mixed, with the former analysis suggesting primarily *Streptococcus* contributions, while the latter identified greatest similarity with *Lactobacillus murinus* and *Candidatus* Arthromitus trajectories (Additional File 8: Figure S4). The differences in the taxonomic composition of the reference data underlying these two approaches could account for these inconsistencies, as could the fact that the KEGG analysis relies on amino acid comparisons while the species trajectories are determined through nucleotide comparisons. Thus, an uncharacterized lipooligosaccharide transporter encoded in *Streptococcus* and other gut microbes decreases in abundance over time at a significantly faster rate in DNR compared to WT mice, starting early in life before weight loss and immune activation.

Type III secretion system (M00332) differed in its temporal change between the lines uniquely before disease onset. Specifically, the module decreased in abundance in WT mice over weeks 1-7, with KO K03225 primarily driving this effect. On the other hand, this module was relatively stable in DNR mice prior to disease onset, and several of the KOs that comprise the module increased in DNR mice in the later weeks (Fig. 4). The discovery of stable, rather than decreasing, abundance of K03225 as an early indicator of IBD in DNR mice is intriguing because Type III secretion systems are used by pathogens to invade the gut community and alter the gut environment (68, 69).

We next examined baseline differences in module abundance between DNR and WT mice at weaning. Early differences could result from genotype-specific selection of the gut microbiome or cage effects. Our models revealed 17 modules with significantly different intercepts (q<0.05), which indicates differences in abundance between the two lines at week 4 (Additional File 9: Table S4). Eight of these modules, including several methanogenesis associated pathways, had positive intercept coefficients, meaning that they were more abundant in DNR compared to WT mice at week 4. Lipopolysaccharide biosynthesis and eight other modules showed the opposite effect and were higher in WT mice at weaning. This early-life variation in the microbiome supports hypotheses that pre-adolescent development of the microbiome can affect health state later in life. However, these temporal relationships are complex: later changes in abundance, as captured by the time by cohort interaction, could reverse the pattern seen at weaning.

To explore temporal dynamics of specific gut taxa, we applied the same GLMM analysis to species abundances. This analysis yielded no significant results at FDR < 0.05, likely due to not having the advantage of grouping components across a higher order variable. While species could be grouped into higher taxonomic entities, the model assumption that members of the same group tend to covary across samples and over time may be violated because members of the same taxonomy may compete or ecologically exclude one another (70). We evaluated this possibility by applying a non-parametric decomposition of variance components (71) to assess whether within-module or within-genus dispersion decomposition patterns were significantly different from those obtained from random permutations of the underlying data. This auxiliary analysis finds that components of functional groups covary more than random while components of taxonomic groups do not (Additional Data File 10: Figure S5). This observation indicates that grouping taxa would violate GLMM model assumptions. Consequently, we instead used a goodness-of-fit test based on functional principal components analysis (FPCA), which is less rigid in its assumption of linearity and capable of borrowing information across species due to the representation of abundance trajectories as combinations of eigen-functions derived from the entire dataset. This test identified seven species that significantly differ in their variation over time between the DNR and WT cohorts (Figure 5; Table 1), including greater increases in abundance over time within DNR microbiomes for *E. coli* and four species from the *Bacteroides* genus, which are associated with gut inflammation (10).

**Figure 5.**
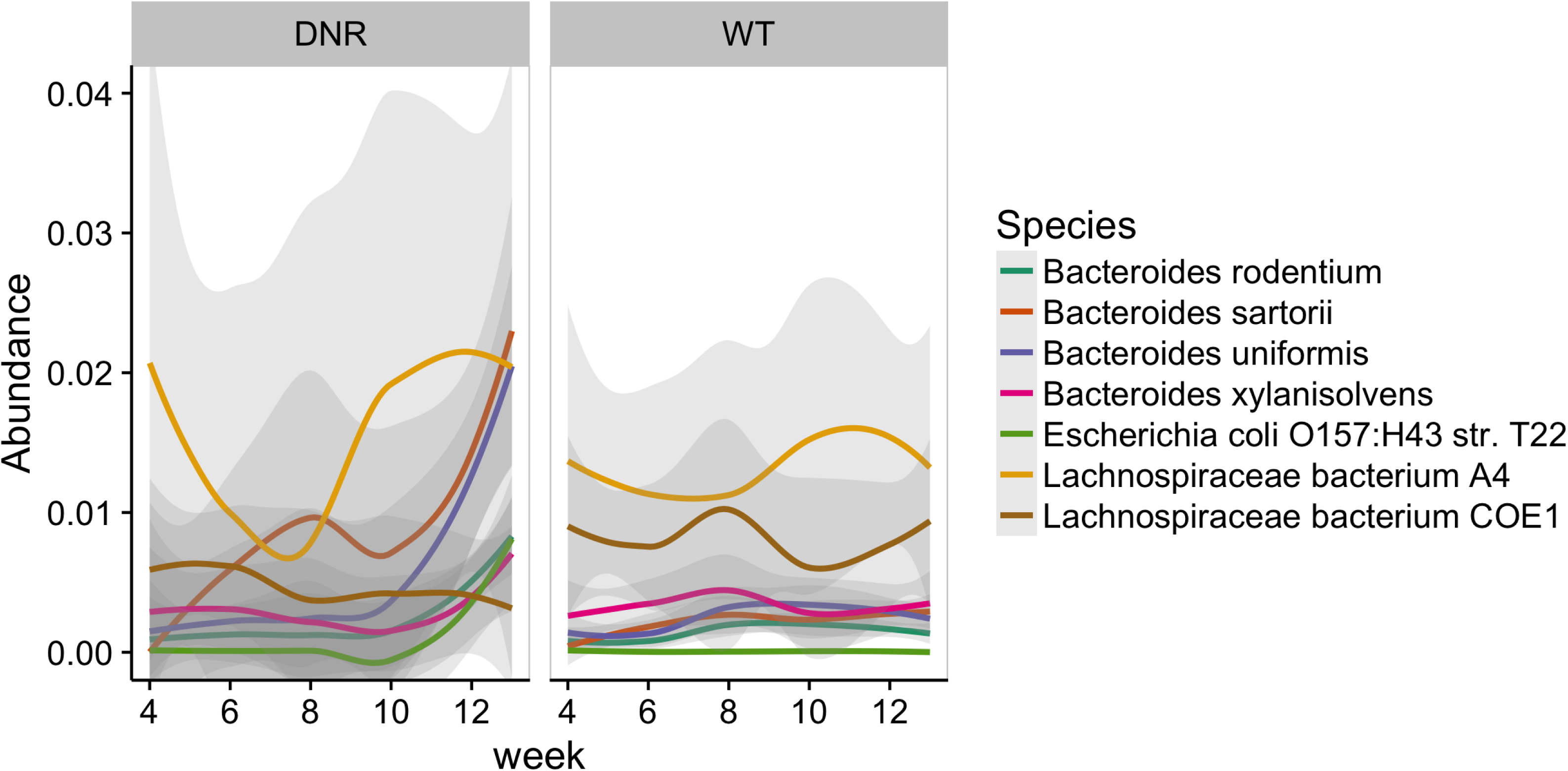
Species that showed significantly different trajectory shapes between DNR and WT groups. These results are based on an FPCA-based goodness-of-fit comparison test that identified 7 species that were different at B-H corrected p-value < 0.05.

**Table 1.** Species with significantly different trajectory shapes in the FPCA-based goodness-of-fit comparisons.

| Species ID | p-value | q-value | Species Name | WT area under LOESS curve | DNR area under LOESS curve |
| --- | --- | --- | --- | --- | --- |
| 54642 | 0 | 0 | *Bacteroides sartorii* | 0.01992 | 0.07699 |
| 57185 | 0 | 0 | *Bacteroides xylanisolvens* | 0.03051 | 0.02506 |
| 57318 | 0 | 0 | *Bacteroides uniformis* | 0.02297 | 0.04506 |
| 58110 | 0 | 0 | *Escherichia coli*<br>*O157:H43 str. T22* | 5.35E-04 | 0.007523 |
| 59684 | 0.0001 | 0.0025 | *Lachnospiraceae*<br>*bacterium COE1* | 0.07203 | 0.04213 |
| 59708 | 0 | 0 | *Bacteroides rodentium* | 0.0136 | 0.01986 |
| 61442 | 0.0013 | 0.02786 | *Lachnospiraceae*<br>*bacterium A4* | 0.119 | 0.1348 |

## Discussion

This study represents the first shotgun metagenomic characterization of IBD development. By using a controlled mouse model, a longitudinal study design, and statistical modeling, we identified novel microbial biomarkers associated with IBD onset and progression. Many of the taxa and functions we implicated have known roles in immune regulation and pathogenicity, making them plausible candidates for stimulating the disease process, while others likely represent responses of the microbiota to changes in host physiology. Ordination and GLMM analyses enabled us to distinguish these scenarios by identifying significant differences between DNR and WT mice over time from weaning through severe disease. We discovered that lipooligosaccharide transport and type III secretion protein abundance trajectories between weaning and immune activation differentiate DNR mice prior to immune activation, making them promising early biomarkers and consistent with a potentially causal role in IBD. Abundances of 17 modules are altered in DNR mice at weaning and could predict IBD risk if they generalize to other mouse models and human disease (see below). Many other modules as well as a few species have altered abundances in DNR mice in later, more severe stages of disease. Functional and taxonomic diversity also show temporal differences in DNR mice that correlate with immune profiles and/or disease progression. Most of these discoveries would have been missed in a cross-sectional study because the disease association is a longitudinal trend.

By using shotgun metagenomics, we were able to investigate both taxonomic and functional characteristics of the IBD microbiome. Both types of data consistently showed differences between DNR and WT mice. For example, beta-diversity analyses revealed increasing divergence of both taxonomic and functional profiles between DNR and WT microbiomes over the last four weeks of the study. In addition, the individual taxa and modules with genotype-specific trajectories predominantly had increased abundance in DNR mice after disease onset. These similarities in the successional diversification of species and genes support the idea that taxonomic changes in IBD have functional consequences that are linked to immune activation. Despite such parallels, our taxonomic and functional results differed in several important ways. Notably, a smaller number of species stratify lines over time as compared to KEGG modules. Furthermore, most of the IBD-associated modules we discovered were not represented solely in singular species and would have been missed by considering information from taxonomic analyses only. These results could be due to disease-associated functional redundancy, wherein a gene that is enriched in DNRs might derive from different species in each mouse. Other potential reasons include (1) higher power due to grouping protein families into modules, and (2) missed taxonomic associations due to the relatively small number of laboratory mouse associated microbes in the genome tree of life (72). Future work should explore how taxa missed by reference-based quantification vary in association with IBD in DNR mice.

Despite finding relatively few species that distinguish DNR mice, we can gain insight into the disease process from what is known about how these taxa interact with the host. It is striking that four of the seven species that change in abundance as IBD develops belong to the genus *Bacteroides,* three of which are more abundant in DNR mice. Several studies have implicated *Bacteroides* in intestinal inflammation. For example, a subset of *B. fragilis* strains carry a proinflammatory metalloprotease toxin that has been identified in 19.3% of patients with active IBD (73), and the inoculation of animals with such strains is associated with severe colitis (10, 74). Subsequent research showed that multiple commensal species of *Bacteroides* could be incorporated into the gut microbiomes of IBD-susceptible genotypes of mice to induce IBD, including mice with TGF-β susceptibility loci (75). Supporting the idea that *Bacteroides* contribute to IBD, we observe a modest increase (q=0.1898) in the hemophore/metalloprotease transport system module (M00328) in DNR mice as disease progresses. These and other mechanistic hypotheses must be tested, because the species of *Bacteroides* we identified are diverse and species within the same genus can exhibit discordant patterns of interaction with host physiology (76).

Cross-sectional and mechanistic investigations of IBD support our finding that disease development is linked to microbiome taxonomy and function (4, 77, 78). The progressive divergence of DNR and WT microbiomes as IBD worsens is consistent with a 16S-based study using a different mouse model of IBD in which gut microbes and imputed functions changed in association with disease status and therapeutically induced remission (22). Additionally, studies in germ-free mouse models of IBD implicate the gut microbiome in disease development. For example, interleukin (IL)-10-knockout mice grown under germ-free conditions do not develop colitis, while conventionally raised mice do (79). Similar findings have been reported for the TRUC mouse model (80). Furthermore, IL-10-knockout (81) and IL-2-deficient (82) mice manifest differential severity of colitis dependent on the types of taxa that colonize their gut. Human studies of IBD have yet to investigate the disease longitudinally. However, our results are consistent with microbiome case-control studies that found significant differences in the taxonomic (11, 83-87) and functional (13, 16, 22) profiles of IBD patients compared to healthy controls, especially in Crohn’s disease. Additionally, clinical administration of antibiotics shows promise for reducing the intestinal inflammation associated with IBD (88, 89). The longitudinal biomarkers we identified are promising new candidates to investigate in the context of human disease onset and progression.

Our analyses identified several modules that implicate a pathogenic effect by the DNR microbiome. For example, DNR mouse microbiomes increase in the abundance of adhesion protein transport modules (M00330) in association with disease, which may help pathobiotic members of the microbiome associate with and metabolize intestinal mucosa (90). Correspondingly, keratan (M00079) and dermatan (M00076) sulfate degradation pathways increase in abundance as disease progresses. Keratan sulfate and dermatan sulfate are glycosaminoglycans (GAGs) that are integral to intestinal mucosa and regulate the permeability of the gut epithelium. These sulfated GAGs are depleted in IBD patients (91), and their metabolism by intestinal bacteria, including *Bacteroides thetaiotaomicron*, contributes to intestinal colonization (92, 93). Furthermore, Crohn’s metagenomes exhibit an increase in GAG degradation pathways (94). DNR guts also have elevated levels of Type III and Type IV secretion systems, which pathogenic organisms leverage to successfully invade the gut microbiome and induce preferable ecological conditions within the gut (68, 69). Curiously, type III secretion abundance shows the opposite effect before immune activation (weeks 4-7), perhaps because of broad shifts in community composition after week 7 or alternatively due to microbes with type III secretion systems invading the LP and becoming less abundant in stool over time. Finally, we observe an increase in modules associated with the biosynthesis of isoprenoids, which have been linked to the stimulation of the mammalian immune system (95). Together these DNR-associated pathways support a pathogenic role of gut microbes in IBD development. Future studies that seek to determine the existence of a microbiome-mediated etiology for IBD should consider these potential mechanisms of disease activation.

Our identification of pathways that change in association with IBD development generates many novel hypotheses about the mechanisms through which gut microbes contribute or respond to disease development. Future studies can explicitly test these hypotheses to discern the cause and effect relationship between the gut microbiome and inflammatory bowel disease. Several KEGG modules with different abundance dynamics in DNR versus WT mice appear to be associated with the microbiome’s acclimation to the disease environment. For example, we observe increases in two-component systems (M00511, M00482) that may contribute to a cell’s ability to manage the elevated oxidative stress that exists during active IBD (96). We also observe increases in pathways associated with cellular chemotaxis (M00515, M00507). This result is consistent with observations of increased cell motility pathways in the gut microbiomes of TRUC mice suffering active colitis using imputation from 16S data (22). This result also aligns with prior work that implicated toll-like receptor recognition of flagellar bacterial antigens in the development of intestinal inflammation (97, 98). Based on these observations, we speculate that chemotaxis pathways help microbiota scavenge the metabolic resources required to survive inside of an inflamed gut or invade the host given that intestinal permeability frequently increases during IBD flare-ups (99).

We also observe several biosynthetic modules that increase in association with IBD development. For example, modules related to the biosynthesis of uridine monophosphate, leucine, proline, and ammonia change in association with disease. These results may suggest that the metabolic preferences and needs of the organisms that comprise the microbiome change as disease develops. Alternatively, it may be that more T cells are entering the gut, becoming activated, and consequently consuming the local resources, which in turn results in bacteria activating biosynthetic pathways to survive and compete. Our finding that pathways associated with ammonia production (M00531) increase in DNR mice is noteworthy because prior studies have found that IBD associates with a lower pH in the intestinal lumen (100), and the production of ammonia by bacteria may serve to buffer such pH changes. Additionally, these pathways are utilized when bacteria metabolize proteins, amino acids, and urea, and the increase in this pathway may indicate a preferential utilization of these substrates by the microbiome or, as above, immune cells, during disease.

Furthermore, we observe increases in modules associated with choline metabolism, specifically betaine and phosphatidylcholine biosynthesis. Recent work has connected the gut microbiome’s production of these metabolites to increased cardiovascular disease risk (101). Our finding is important because a growing number of studies indicate that IBD patients have an increased risk of developing cardiovascular disease, especially during flare-ups (102, 103). The mechanisms underlying this increased risk are not well resolved, but may relate to a proposed explanation for the increased cardiovascular disease risk observed in HIV-infected patients (104, 105). Under this model, changes in the relative proportion of protective to pathobiotic gut microbiota, especially those capable of translocating across the gut epithelium, activate a chronic systemic inflammation that increases cardiovascular disease risk. It is thus tempting to speculate that, based on our observations in these mouse models of disease, IBD and perhaps HIV-associated changes in microbial metabolism of choline contribute to or at least indicate this increased risk of cardiovascular disease.

Another intriguing hypothesis emerges from our observation that heme transport genes are elevated in DNR mice as IBD develops. Bacteria use this module to scavenge iron from the environment. Iron is a crucial component for many cellular processes, but gut microbes seldom have access to free iron and instead sequester it from host sources, such as heme (106, 107). Heme concentrations may be increased in IBD, as a common feature of the disease is intestinal bleeding (108). Hence, we hypothesize that gut microbes that can take advantage of this heme may flourish in DNR mice. It is intriguing to further speculate that microbial sequestration of heme contributes to IBD (e.g., through signaling to the immune system) or to iron deficiency in IBD patients (109).

One surprising discovery was an increase in pathways associated with the production of benzoate (M00538) in DNR mice. Benzoate is a carboxylic acid produced by microbial degradation of dietary aromatic compounds and is a precursor of hippurate biosynthesis in mammals (110). Prior work suggested that hippurate may be a useful diagnostic of Crohn’s disease given that it is found at significantly lower levels in the urine of patients (110) and that the gut microbiome’s production of benzoate is responsible for these differences in urinary hippurate (111). Our results are inconsistent with this prior work in such that they indicate that intestinal benzoate biosynthesis is higher in sick animals. This difference may be due to variation in the host species being investigated, including how benzoate is subsequently metabolized in the gut or by the host. Alternatively, the potential of the DNR microbiome to make excess hippurate may not be realized given that we performed DNA sequencing. Future mechanistic studies could measure benzoate and hippurate and quantify the benzoate proteins at the RNA or protein level in DNR versus WT mice.

Most of the taxonomic and functional IBD biomarkers we identified are increasingly abundant in DNR mice throughout the disease process. But three modules show the opposite trajectory and decrease in abundance over time in DNR relative to WT mice: melatonin biosynthesis (M00037), lysine biosynthesis (M00031), and lipooligosaccharide transport (M00252). Melatonin has a dual effect on the immune system, acting in a stimulatory manner in early infection, and in an immunomodulatory manner in cases of prolonged inflammation (112). The effects of melatonin produced by gut commensals have not been studied as extensively as those of endogenous melatonin. Traditionally, melatonin acts as a potent antioxidant, although additional quorum signaling functions in bacteria have been recently reported (113). The reduction in melatonin biosynthesis capacity observed in the DNR mice could be caused by the expansion of species that can tolerate a highly oxidative environment (114) or microbes that utilize other strategies for neutralizing reactive oxygen species. Without metabolite data, it is not possible to definitively say that the final concentrations of melatonin are reduced in the disease state, since the decrease can be offset by host production. With respect to lysine biosynthesis, this module is also depleted in human IBD microbiomes (115), indicating that there may exist similar mechanisms of interaction between disease context and the gut microbiome across species. Future work should empirically test the potential role of these microbiome functions on the development of IBD, especially in individuals that are genetically susceptible for the disease.

Lipooligosaccharide transport is the only module to show significant differences in abundance trajectories both pre- and post-activation. Intriguingly, it is consistently lower in DNR versus WT mice throughout our study with the largest difference during weeks 4-7, prior to immune activation and disease symptoms. This finding initially seems surprising, because lipooligosaccharides are the major glycolipids that are produced by mucosal Gram-negative bacteria and are known to have proinflammatory effects (116). However, the two genes (NodI and NodJ) in the lipooligosaccharide transport system are present across diverse prokaryotes, and the substrates of this two-component ABC transporter are not characterized beyond lipo-chitin oligosaccharide export in rhizobial bacteria (117, 118). Determining what this system transports in the mammalian gut and how its function changes in IBD is an exciting future direction. Regardless of mechanism, the consistent and pre-symptomatic depletion of lipooligosaccharide transport genes in DNR mice make this module a promising candidate biomarker for predicting and diagnosing IBD.

We relied on a mouse model to quantify the longitudinal interaction between the gut microbiome and disease because the extensive inter-individual variation in human genetics, lifestyle, microbiome composition, and disease status and severity can complicate study design, analysis, and interpretation. We used the DNR mouse model because it is relevant to our understanding of the mucosal immunological dysregulation that occurs during human IBD and, consequently, its interaction with the gut microbiome. Indeed, we observe immune activation in the blood of the DNR mice that is consistent with what has been observed in human IBD (119). The phenotype observed in DNR mice is akin to severe Crohn’s disease with relatively substantial immunological activation and weight loss by week 12. Interpretations of the microbiome-disease interaction in this model should be considerate of this relatively severe disease status. Alternative mouse lines may be better models for other forms of IBD. Another consideration is that we found some baseline differences in microbiome protein abundances in DNR mice at weaning that may be specific to this genetic model of IBD. Ultimately, comparisons between our results and those obtained by the integrated Human Microbiome Project (iHMP) (120) which is longitudinally evaluating the microbiome and immune status of IBD patients, will clarify the relevance of the findings produced by the DNR model to human populations. Additionally, future research should use this model and build upon our findings to clarify how TGF-β induced differentiation and function of T cells interacts with the taxonomic structure and function of the gut microbiome.

Overall, our results indicate that the development of IBD is associated with corresponding changes in the operation of the gut microbiome. Microbial taxa and KEGG module abundances vary over time and in association with immune activation. Furthermore, our results suggest that the gut microbiome may contribute to disease by activating inflammation through metabolism of mucosa and by expressing proinflammatory and downregulating anti-inflammatory metabolites. Because our study relied on the imputation of microbiome function from DNA sequences, we cannot definitively conclude that the observed differences in the microbiome’s functional profiles manifest as differences in the metabolites produced by the microbiome. Future research that applies direct measurements of microbiome function should be used to validate and expand the results presented here. Regardless, our results hold promise for our understanding of microbiome-mediated IBD disease mechanisms and the potential of using microbiome sequencing of patient stool to classify and potentially even predict disease.

## Methods

### Growth of mice and microbiome sampling

We bred two cohorts of DNR and WT littermate control animals in the Gladstone Institutes mouse facility as follows. CD4-dnTβRII (DNR) animals were crossed to RAG1-/- background to eliminate the T cell mediated IBD, and were transferred from Yale University to Gladstone Institutes in 2010. To initiate experiments described in this study, DNR-RAG1-/- males were bred with C57BL/6N female animals, and DNR-RAG1-/+ progeny were again crossed to C57BL/6N females to generate a combination of RAG1-/+ and RAG1+/+ DNR and WT age-matched littermate controls. Animals were given regular chow consisting of irradiated PicoLab Rodent Diet 20 (LabDiet). Only female animals were used in this study. Four co-housed WT-RAG1+/+ and five co-housed DNR-RAG1+/+ littermates were followed longitudinally for 15 weeks and fresh fecal samples were collected weekly and stored at -30° C until they were subject to microbiome processing. All mice from both cohorts were weighed weekly. All animal experiments were conducted in accordance with guidelines set by the Institutional Animal Care and Use Committee of the University of California, San Francisco.

### Immune sampling

Tail vein blood samples were collected weekly from a parallel cohort of “bleeder” mice to quantify how their immune status changes over time (n=6 WT, n=6 DNR). These are distinct individuals from the “pooper” mice subjected to stool metagenomics (same colony and time period), in order to prevent repeated tail vein blood sampling from affecting the health or microbiota of the cohort of pooper mice. Specifically, ∼100 µl (2-3 drops) of blood from tail vein was added to 30 µl of 1x heparin (500 units/ml). 500 µl of 1x ACK lysis buffer (Lonza) was added directly to the cells and incubated at room temperature for 2–3 min. Cells were centrifuged at 4000 rpm for 5 min. The top layer was aspirated and another 500 ml of 1x ACK lysis buffer was added followed by centrifugation. Cells were resuspended in FACS buffer (PBS + 0.5% FBS) and after blocking Fc receptors with anti-CD16/CD32, single-cell suspensions were incubated with FITC CD4 (GK1.5), PE CD62L (MEL14), PerCP-Cy5.5 CD8a (53.6.72), and APC CD44 (IM7) mouse antibodies for 30 min at 4°C. Stained cells were washed and acquired on an Accuri C6 cytometer (BD). Blood lymphocytes were gated on CD4+ or CD8+ fractions and percentage of activated/memory (CD44hi) among CD4+ and CD8+ T cells was determined using FlowJo software (Tree Star Inc.). This cohort was separated from those subjected to microbiome sampling to eliminate the effect that repeated bloodletting might have on the microbiome. At 15 weeks of age, two WT and three DNR from the non-bleeding animals were euthanized. Spleen and mesenteric lymph nodes were then processed into single-cell suspension and subjected to ACK lysis and cell surface staining as described for PBMCs. The status of T cell activation was quantified and found to highly correlate with the blood immune status of their “bleeder” littermates (Additional Data File 2: Figure S2 and Additional Data File 3: Table S1).

### Metagenome sequencing and analysis

QIAamp DNA Stool Mini Kits (QIAGEN, Valencia, CA, USA) were used to extract DNA from stool samples collected from weeks 4, 6, 8, 10, and 12. Samples were incubated in an elevated water bath temperature of 95 C to increase the lysis of bacterial cells, as per manufacturer instructions. The MoBio PowerFecal DNA isolation kit (MOBIO, Carlsbad, CA USA) was used as per manufacturer instructions to process stool samples collected at weeks 5 and 13. Kit type was adjusted for in statistical modeling to account for any potential differences in extraction bias between the two methods.

Purified DNA was prepared for shotgun metagenomic sequencing using the Nextera XT library preparation method (ILLUMINA, San Diego, CA USA). Libraries were quality assessed using qPCR and a Bioanalyzer (Agilent Technologies, Palo Alto, CA USA) and subsequently sequenced using an Illumina HiSeq 2000. This produced an average of 74,427,303 100-bp paired-end sequences per sample. Metagenomic reads were quality controlled using the standard operating procedure defined by the Human Microbiome Project Consortium (121) as implemented in shotcleaner (122). Briefly, reads were quality trimmed using prinseq (123) and mapped against the mouse reference genome sequence (GRCm38) using bmtagger (124). Exact duplicate reads were collapsed and the subsequent high-quality data was subject to taxonomic and functional annotation. Functional annotation of metagenomes was conducted using ShotMAP as described in (16) with, Prodigal (125) to call genes and RAPsearch2 (126) to identify metagenomic homologs of the KEGG database (downloaded Feb. 2015). Reads mapping to mammalian sequences in the KEGG database were discarded and the subsequent data was used to quantify the abundance of each KEGG Orthology Group (KO) using the RPKG abundance statistic (127). Metagenomes were taxonomically annotated using MIDAS as described in (72).

### Statistical analyses and modeling

The functional and taxonomic similarity between metagenomic samples was assessed using non-metric multi-dimensional scaling (NMDS) as implemented through the nmds function in the labdsv R package (128). Ordinations were visualized using the ordiplot function in the vegan R package (129). For the functional similarity analysis, the vegdist function from the vegan R package quantified the Bray-Curtis dissimilarity based on KEGG module abundances. The taxonomic analysis used the generalized Unifrac (130) distances (alpha=0.5), which were obtained by using the taxonomic tree from the Living Tree Project (131) and matching the genus and species component of tree leaf labels to the corresponding components of the MIDAS species labels in our data. Assessment of the significance of the clustering of samples in these ordination plots was conducted using PERMANOVA as implemented by the adonis function in R.

The compound Poisson generalized linear mixed effects model implemented in the cplm package in R (132) was used to find KEGG modules with a significantly different time trend between groups while controlling for static differences between the lines and DNA extraction procedure (QIAGEN versus MOBIO). Random intercepts and slopes for both subjects and contributing KOs were used to capture variation between subjects and between genes while focusing on the large scale shifts over the whole collection of abundance profiles contributing to a module. As described more thoroughly in Supplementary Text S1, the general computational procedure consisted of subsetting the data to each module’s relevant KO abundances and fitting a full model that described the RPKG abundance as a function of time, group, time by group interaction, sequencing kit and random effects of each KO and individual. We then use two reduced models, dropping first the interaction term and then the group term, to obtain p-values via likelihood ratio tests. This is one of the recommended significance testing approaches for mixed models since it avoids using approximations for the residual degrees of freedom that would be necessary to test significance via the t-statistic (66). To limit the number of modules tested, the input data were run through the MinPath algorithm (133) to select a parsimonious set of modules based on the KOs present. The union of all samples’ individual parsimonious sets was used as the final set of tested modules. The approach of testing the dynamics of an entire module by fitting a single GLMM to a set of multiple genes’ temporal abundances is modeled on the TcGSA method of Hejblum et al. (134), with the modification of using a different response distribution (the Tweedie compound Poisson). Significant modules were selected at the 0.05 FDR threshold after controlling for multiple testing via the Benjamini-Hochberg procedure (B-H). Species time trend differences were tested with the same approach, minus the grouping of multiple trajectories. Additional details of our modeling approach can be found in Additional Data File 5: Text S1. All of the code used in this analysis is available at the following URL: <u>https://github.com/slyalina/Mouse_IBD_2017_paper_supporting_code</u>.

To differentiate functional changes occurring prior to immune activation, we fit a second hinge regression to the abundances of modules that were found to have a significant time by group interaction in the main GLMM analysis. This second regression placed a break point at week 7, which represents the point at which immune activation initiated (Fig. 1). This allowed for two sets of slopes (before disease onset and after) and two sets of time by group interactions (representing deviations of DNR slopes from WT before-onset and after-onset slopes).

Alterations in the species trajectory curves were additionally tested with an alternate method aimed at highlighting differences in shape rather than slope. This method was an implementation of the FPCA-based difference in goodness-of-fit approach described previously in (135). The permutation-based p-values from this analysis were B-H corrected and species passing the 0.05 FDR threshold were retained.

To test the hypothesis that the distribution of all modules’ between-KO/within-KO dispersion decomposition statistic is significantly different from random when grouping functional trajectories (KOs into modules) but is not significantly different from random when grouping taxonomic trajectories (species into genera) we used the DISCO (71) non-parametric test to obtain the real distributions of the test statistic in the two scenarios, as well the simulated null distributions that arise when generating random groupings of KOs and species. We then performed a Kolmogorov-Smirnov test to compare the true distributions with their simulated counterparts.

## Accession numbers

Metagenomic sequences are available through GenBank accessions SAMN06921515 - SAMN06921563.

## Acknowledgements

We are grateful to Stephen Nayfach for helpful discussions. We also appreciate the services provided by the Gladstone genomics core, UCSF sequencing core, and the Center for Genome Research and Biocomputing.

## Supplemental Files

**Additional File 1: Figure S1.** Longitudinal phenotypic monitoring of T cell activation in PBMC of WT and DNR “bleeder” mice. Representative FACS plots from WT bleeder #89 and DNR bleeder #85 at indicated time points. The gate is set on CD44^hi^ population to show the percent of activated effector and memory T cells over time.

**Additional File 2: Figure S2.** Phenotypic monitoring of CD8 T cell activation in WT and DNR “pooper” mice post disease onset. Representative FACS plots from spleen or mesenteric lymph node (MLN) isolated from WT pooper #90 and DNR pooper #83 at week 15. Fraction of activated (CD44^hi^) and effector (CD44^hi^ KLRG-1^hi^) CD8+ T cells are shown.

**Additional File 3: Table S1.** Project metadata

**Additional File 4: Figure S3.** The functional alpha-diversity of the gut microbiome as measured by Shannon entropy differentially varies over time across cohorts (left). In fact, there is significantly greater variation in the Shannon entropy between lines before and after disease onset (right).

**Additional File 5: Text S1.** Modeling methods

**Additional File 6: Table S2.** KEGG modules that exhibit significant group by time interaction coefficients, indicating that they differentially diversify over time between the two lines.

**Additional File 7: Table S3** - KEGG Modules with significant interactions in the segmented GLMM analysis

**Additional File 8: Figure S4** - Two different analyses potentially explain the taxonomic origins of the lipooligosaccharide transport KOs that were observed. (a) Species identities of KEGG ortholog sequences that recruited reads when generating the relevant KO abundances. (b) Distance covariance values trajectories of K09694 and K09694 and all species, with asterisks marking those dCov values that were significantly non-zero after B-H multiple testing correction.

**Additional File 9: Table S4** – KEGG modules with significant intercepts, indicating that they exhibited significantly different abundances between lines at the initial time point.

**Additional File 10: Figure S5** - Distributions of F-statistics computed by the DISCO method for KO and species vectors within module and genus groupings respectively, compared to values from permuted groupings. Kolmogorov-Smirnov tests show significant difference between real and permuted distributions in the functional groupings, but no significant difference in the taxonomic groupings.

